# Toxic metals increase root hair density by reducing epidermal cell length

**DOI:** 10.64898/2026.02.03.703618

**Authors:** Julia Zheku, Thea Do, M. Arif Ashraf

## Abstract

Root hair cells, which are instrumental in water and nutrient uptake, grow polarly from the epidermal cell layer of the root. Furthermore, plants growing in challenging climates and complex soil environments acclimatize their root hair phenotypes, either by altering root hair length or density. Toxic metal stress is one of the major environmental stresses faced by plant roots. In this study, we demonstrate that toxic metals, such as chromium and arsenite, increase root hair density as an adaptive response. Using the model plant *Arabidopsis thaliana* and other crops plants, like *Zea mays* and *Triticum aestivum*, we further discovered that increased root hair density is caused by shorter epidermal cell length rather than alteration of epidermal cell fate. This study highlights the adaptive cellular and anatomical features of roots during toxic metal stress in evolutionary diverse plant species.

## Introduction

Industrial waste is a major contributor to global environmental pollution, particularly through the release of toxic metals into soil and water systems. Soil contamination with toxic metals such as lead (Pb), chromium (Cr), and arsenite (As) poses a serious threat to plant development, agricultural productivity, and food security (Angon et al. 2024). As the primary interface between plants and soil environment, roots are directly exposed to toxic metal ions. Due to similar physicochemical properties between toxic metals and essential minerals, roots absorb these metals through existing nutrient transport systems (Angulo-Bejarano et al. 2021). For instance, potassium transporters uptake cesium (Ashraf et al. 2021) and phosphorus efflux carriers transport arsenic (Ashraf et al. 2020) due to chemical similarity. Once taken up these toxic metals accumulate in edible plant tissues and ultimately enter the food chain, causing irreparable harm to human health (Zhao et al. 2024).

Within the root system, the epidermis or the outermost cell layer is characterized by root hairs, which are tubular, unicellular extensions of specialized epidermal cells (Schiefelbein et al. 2009; Salazar-Henao et al. 2016; Orr and Ashraf 2025). These cells play, above all, a cricitcal role in increasing the root surface area but additionally support water and nutrient uptake from the soil but also aid in water retention, facilitate root anchorage, and maintain balanced interactions with rhizosphere microorganisms (Schiefelbein et al. 2009; Salazar-Henao et al. 2016; Burak et al. 2021; Orr and Ashraf 2025). As a result, plants use root hair development as an adaptive mechanism by increasing or decreasing root hair length and density. For instance, low temperature stress increases the root hair length in maize and Arabidopsis (Zhou et al. 2024; UrzÚa Lehuedé et al. 2025). Conversely, in phosphorus poor soil, plant roots increase the root hair density as an adaptive mechanism (Wendrich et al. 2020).

Toxic metals, such as cadmium and arsenic, are known to enhance root hair density (Bahmani et al. 2016; Kohanová et al. 2018). The fundamental cellular mechanism of high root hair density as a result of toxic metal exposure is still elusive. In this study, we took advantage of chromium and arsenite to induce high root hair density in *Arabidopsis thaliana, Zea mays*, and *Triticum aestivum*. Using root hair cell specific fate markers and quantitative live cell imaging, we have discovered that toxic metals reduce epidermal cell length, which ultimately enhances root hair density. This mechanism is fundamentally important to advance our understanding about increased root hair density as an adaptive mechanism for roots in toxic soil environments.

## Results

### Chromium (Cr) and arsenite (As) induce high root hair density in Arabidopsis, maize, and wheat

Toxic metals, such as lead (Pb), cadmium (Cd), chromium (Cr), and arsenite (As), inhibit the primary root growth of Arabidopsis (Hazelwood et al. 2025a). Previous studies suggested that Cr and As increase root hair density (Bahmani et al. 2016; Kohanová et al. 2018). Both of these studies germinated seeds on Cr and As containing plates and observed the phenotype either after 7 days or 14 days (Bahmani et al. 2016; Kohanová et al. 2018). As a result, it is hard to deduce whether increased root hair density is due to toxic metals or a secondary effect. To resolve this issue, we transferred the 3-day old light-grown wildtype seedlings to agar plates supplemented without (control) and with 10µM Cr or 10µM As (Figure 1A). We decided on a 10µM Cr and 10µM As concentration as well as a 2-day incubation time based on our previous studies regarding primary root growth (Ashraf et al. 2020; Hazelwood et al. 2025a). We initially observed the inhibition of primary root growth, which was consistent with our previous findings, as well as a higher root hair density in presence of 10µM Cr and 10µM As (Figure 1A). Since the increase in root hair density developed within shorter incubation time (2 days), we confirm that this is an early adaptive response during toxic metal stress.

**Figure 1.**
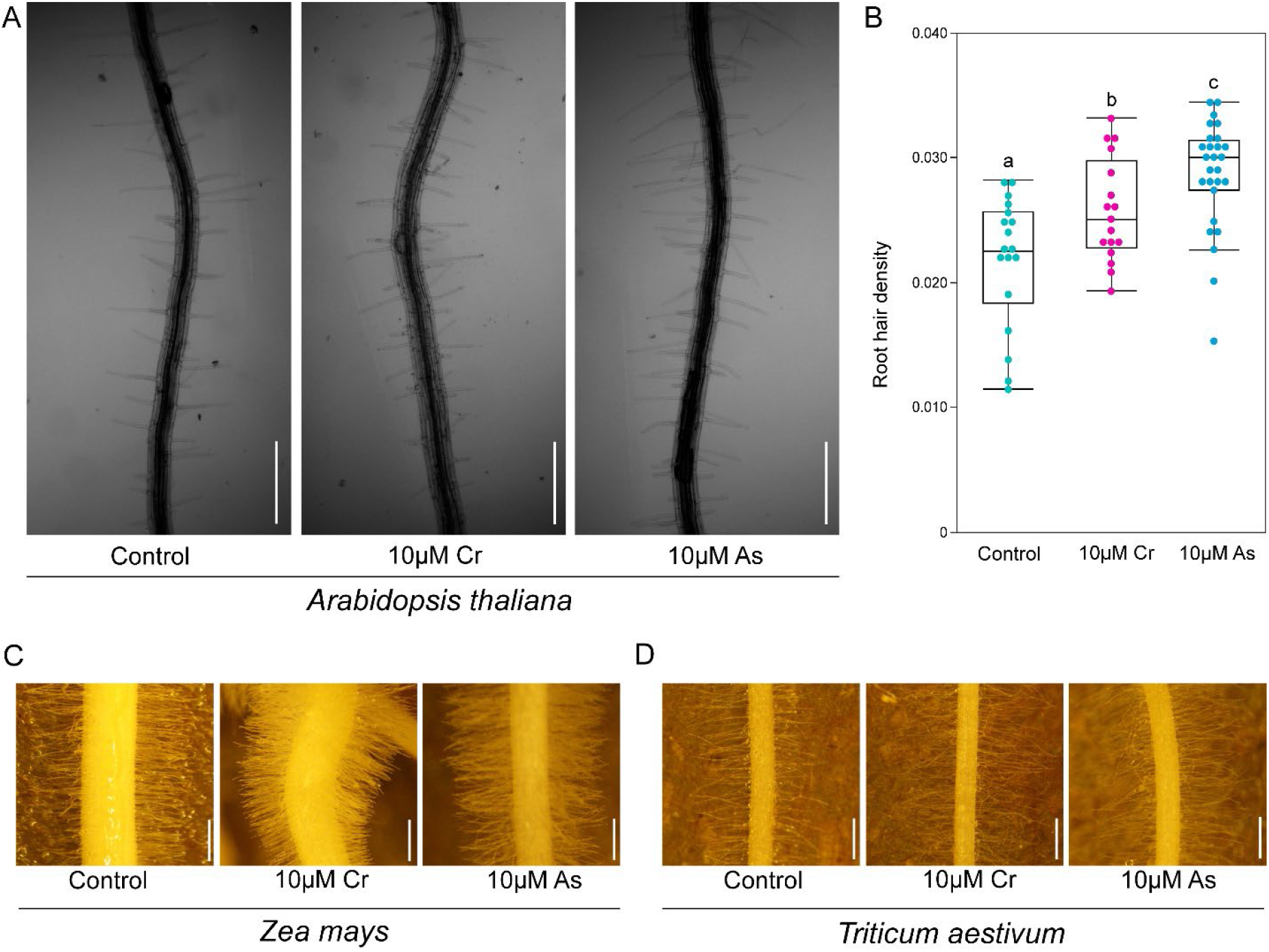
Effects of chromium (Cr) and arsenite (As) on root hair density in Arabidopsis, maize, and wheat. (A) Representative images of Col-0 root hair density under control, 10μM Cr and 10μM As treatments. Scale = 100μm. (B) Quantification of the root hair densities from (A). Statistical test is performed based on Tukey’s Honest test. Groups labeled with the same letter are not statistically different from each other (alpha = 0.05). Boxplots show median values (center line), 25th to 75th interquartile range (box) and 1.5*interquartile range (whiskers). Root hair density phenotype of maize (C) and wheat (D) in absence (Control) and presence of 10μM Cr and 10μM As. Representative images taken over the course of a 2-day exposure. Scale= 1cm.

Furthermore, we quantified the root hair density (number of root hairs divided by the area) and found that there is an ∼1.25 and ∼1.5 times increase in root hair density in the presence of 10µM Cr and 10µM As, respectively (Figure 1B). Previous study observed a 39% increase of root hair density in presence of 10µM As and after 7 days incubation (Bahmani et al. 2016). Our study discovered a ∼25% increase of root hair density in presence of 10µM As after 2 days of incubation, further confirming this trait is an early adaptive response that was previously investigated over longer time period.

Additionally, we observed that among control, 10µM Cr and 10µM As conditions, root hair density is a gradient and varies (root hair density: As > Cr > Control). This is an ideal set up to further explore the cellular and molecular mechanism of toxic metal-induced increased root hair density.

We hypothesized that if increased root hair density is an early adaptive response to toxic metal stress, crop plants’ root systems will respond similarly to Arabidopsis roots. We tested both maize and wheat root in presence of 10µM Cr and 10µM As conditions during the early root development (Figure 1C,D). Interestingly, there is a significant increase of root hair density for maize and wheat root in presence of 10µM Cr and 10µM As over a 2-day long exposure(Figure 1C,D). Altogether, our results suggest that higher root hair density is an early adaptive response during toxic metal stress across diverse plant species.

### Root hair cell specific fate markers remain same during chromium (Cr) and arsenite (As) stress in Arabidopsis

Increasedroot hair density can be caused by alteration of root hair cell fate. For instance, the genes GL2, WER, and TTG express in non-root hair cells and the corresponding mutants (*gl2, wer*, and *ttg*) demonstrate a higher root hair density (Galway et al. 1994; Masucci et al. 1996; Lee and Schiefelbein 1999). We tested non-hair cell type specific markers *proGL2::GUS* and *proCPC::GUS*; and hair cell specific marker *EXP7::NTF* in the presence of 10µM Cr and 10µM As (Figure 2). Surprisingly, we did not observe any alteration of either hair or non-hair cell fate markers after 2-days of incubation at 10µM Cr and 10µM As (Figure 2). Among these cell type specific marker lines, we visualized *EXP7::NTF* with propidium iodide staining. As a result, we not only observed the hair cell specificity, but also the epidermal cell length.

**Figure 2.**
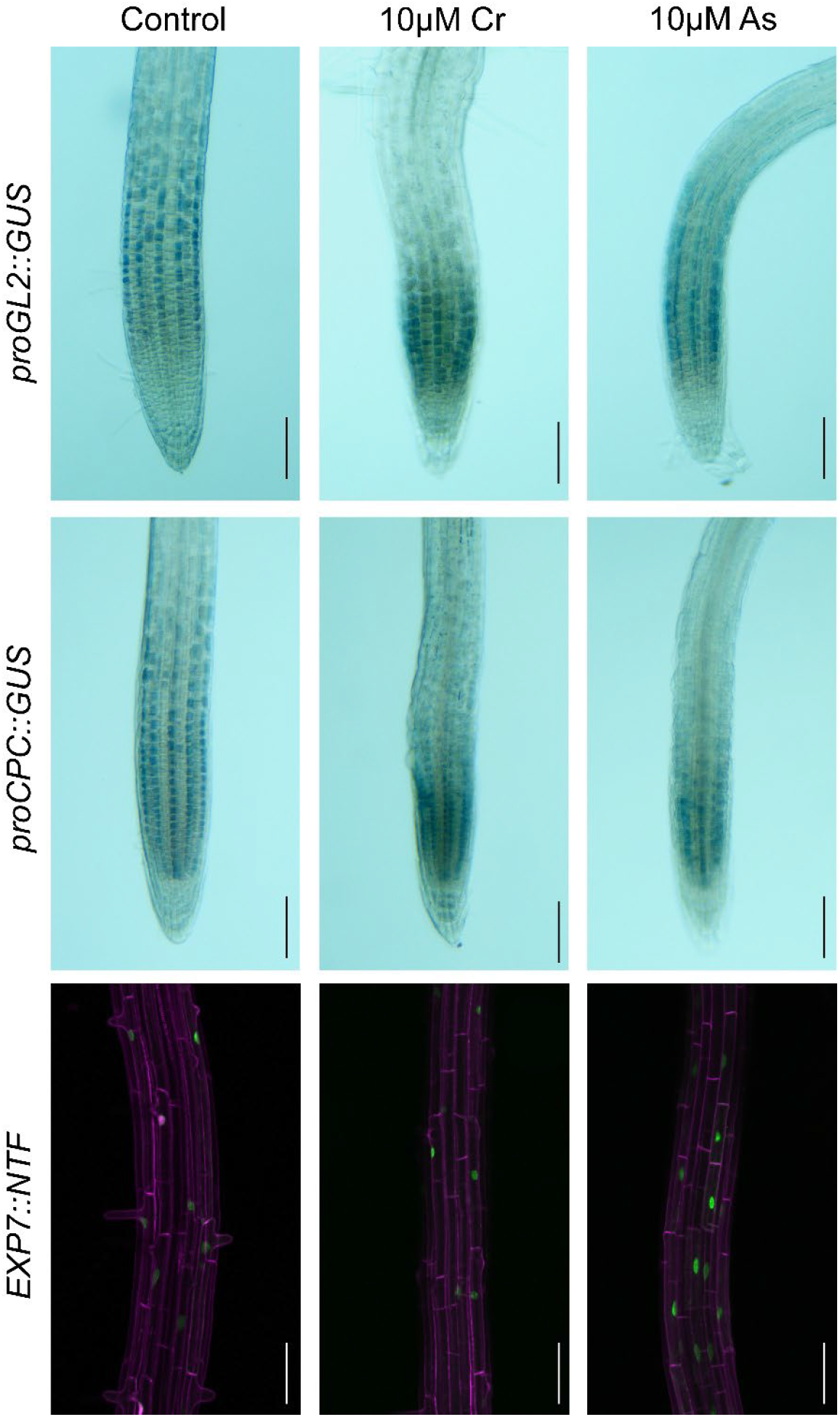
Visualization of non-hair and hair cell specific fate markers in presence of chromium (Cr) and arsenite (As) in Arabidopsis. *proGL2::GUS* (top panel), *proCPC::GUS* (middle panel), and *EXP7::NTF* (right panel) in absence (Control) and presence of 10μM Cr and 10μM As. Scale = 100μm.

Interestingly, we observed that epidermal cell length is shorter in presence of 10µM Cr and 10µM As (Figure 2). These data suggest that the higher root hair density is probably caused by shorter epidermal cell length and not alteration of hair or non-hair cell fate.

### Epidermal cell elongation is reduced during chromium (Cr) and arsenite (As) stress in Arabidopsis

To investigate the epidermal cell length, we utilized plasma membrane localized LTI6b-EGFP in absence (Control) and presence of 10µM Cr and 10µM As (Figure 3A). We observed a sharp reduction in epidermal cell length due to 10µM Cr and 10µM As treatments. Our quantification further confirms this observation (Figure 3B).

**Figure 3.**
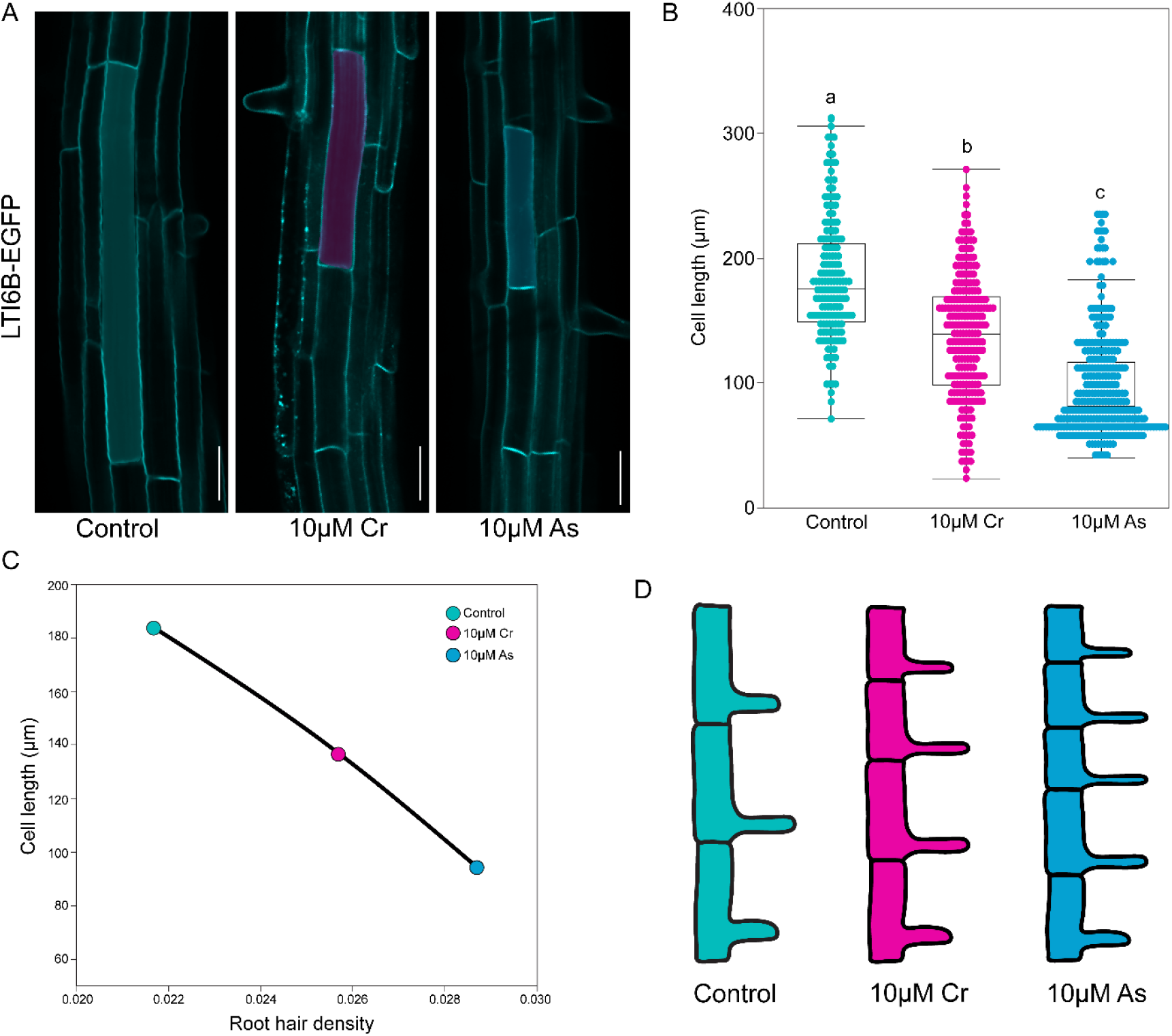
Effects of chromium (Cr) and arsenite (As) on root epidermal cell length in Arabidopsis. (A) Representative confocal microscopy images of root epidermal cells, visualized by LTI6b-EGFP, from control, 10uM Cr, and 10uM As treatments. Scale = 50μm. (B) Quantification of cell lengths from (A). (C) Correlation between root hair density and epidermal cell length from control, 10uM Cr, and 10uM As conditions. Statistical test is performed based on Tukey’s Honest test. Groups labeled with the same letter are not statistically different from each other (alpha = 0.05). Boxplots show median values (center line), 25th to 75th interquartile range (box) and 1.5*interquartile range (whiskers). (D) Graphical model demonstrating the relationship between epidermal cell length and root hair density.

Furthermore, between Cr and As, the effect on epidermal cell length is more severe in As compared to Cr (Figure 3B). This quantitative data of epidermal cell length (epidermal cell length: As < Cr < Control) (Figure 3B) is reminiscent of the root hair density (root hair density: As > Cr > Control) (Figure 1B) data from our initial observation. Interestingly, the root hair density perfectly correlates with the epidermal cell length from the control, 10µM Cr, and 10µM As conditions (Figure 3C). Altogether, our study suggests that toxic metal-induced higher root hair density is caused by the reduction of epidermal cell length (Figure 3D).

## Discussion

The outermost cell layer of root, or epidermis, faces direct contact with the soil and must respond to environmental anomalies. Root hairs are polar outgrowth of epidermal cells and their developmental and genetic circuits are controlled at the cellular level (Schiefelbein et al. 2009; Salazar-Henao et al. 2016; Orr and Ashraf 2025). As a consequence, adaptive mechanisms, such as higher root hair density during toxic metal stress, are important to understand at the cellular level as well as in evolutionary diverse root systems.

Previous studies highlighted that higher root hair density is a consequence of genetic alteration of cell fate (Galway et al. 1994; Masucci et al. 1996; Lee and Schiefelbein 1999; Bahmani et al. 2016; Kohanová et al. 2018)The majority of these studies ignored the parameter of epidermal cell length. Interestingly, epidermal cell length alone can contribute to higher root hair density. For instance, the conditional wheat mutant, *stumpy*, demonstrates higher root hair density due to shorter epidermal cell length in the presence of excess calcium (Zeng et al. 2024). Our study provides a similar mechanism, where the epidermal cell length alone controls the root hair density as an adaptive mechanism during toxic metal stress (Figure 3D).

The root hair and non-hair cell fate markers used in this study are not readily available in other plant species besides Arabidopsis. As a result, it is difficult to track the early hair and non-hair cell fate in non-model grasses and crop plants. In contrast to cell fate marker lines, epidermal cell imaging is easily accessible to study adaptive environmental stress responses in any root system. In our study, we utilized a range of species, Arabidopsis, maize, and wheat, which depict a diverse evolutionary history.

The eudicot Arabidopsis is separate from monocot grasses by 160 years of evolution (Kumar et al. 2017). Within the group of monocot grasses, maize and wheat are separated by 49 million years of evolution (Kumar et al. 2017). Overall, the fundamental discoveries from this study are widely applicable to a range of environmental stresses and evolutionary diverse root systems.

## Methods and materials

### Plant materials

*Arabidopsis thaliana* Col-0 (as wild-type), *proGL2::GUS* (ABRC stock: CS8851), *proCPC::GUS* (ABRC stock: CS6497), *EXP7::NTF* (ABRC stock: CS2106179), LTI6b-EGFP (Kurup et al. 2005), Wheat (variety *Ladoga*, Prairie Garden Seeds; https://prairiegardenseeds.ca/) and Maize (B73 from MaizeGDB; https://www.maizegdb.org/) were used in this study.

### Plant growth condition

Arabidopsis seeds were sterilized with 1mL 70% ethanol for 10 minutes, washed twice with 1mL autoclaved ultrapure water inside the laminar flow hood, and placed on the surface of ½ Murashige and Skoog (MS) media (BioWorld; Cat.# 30630058) containing 1% sucrose (BioWorld; Irving, TX) and 1% agar (BioWorld) in square petri dishes (Simport; Bernard-Pilon Beloeil, Canada). Dish edges were secured with ½ inch micropore tape (Amazon; Seattle, WA), covered with aluminum foil, and kept at 4°C for 2 days. After 2 days, plates were bound with rubber bands and placed vertically at 23°C with a 24h light cycle, as described before (Hazelwood et al. 2025b, 2025a). 3-day-old seedlings were transferred to plates with 10uM Cr or 10uM As added to the agar-based media or with (control).

Before experimental use, wheat and maize were first sprouted on square petri dishes (Cat #26-275; Genesee Scientific) then transferred and subjected to treatments. Seeds were put onto dishes lined with a moist paper towel, covered with aluminum foil, and placed at 4°C for 2 days. After 2 days, the dishes were taken out from 4°C and arranged flat at 23°C with a 24h light cycle, allowing the seeds to grow for 2 days before transfer. The germinated seedlings were placed into plastic boxes (16.5 x 8.5cm; Temu; https://www.temu.com/) also lined with a moist paper towel and maintained for 2 days afterward at 23°C. The seedlings were watered daily either with sterilized water (control) or with solutions containing 10µM Cr or 10µM As. Wheat and maize were imaged for root hair phenotypes over the course of the 2-day treatment.

### Chemical treatment

Potassium chromate (Sigma Aldrich; St. Louis, MO) and Sodium (meta)arsenite (Sigma Aldrich; St. Louis, MO) were dissolved in water to prepare the stock solution. For treating seedlings, 10μM Cr and 10μM As final concentrations were obtained by mixing with either ½ Murashige and Skoog (MS) media for Arabidopsis or water for wheat and maize.

### GUS staining

GUS staining was performed based on a previously described method (Ashraf and Rahman 2019; Hazelwood et al. 2025b, 2025a). *proGL2::GUS* and *proCPC::GUS* seedlings were transferred to GUS staining buffer (100 mM sodium phosphate, pH 7.0, 10 mM EDTA, 0.5 mM potassium ferricyanide, 0.5 mM potassium ferrocyanide and 0.1% Triton X-100) containing 1mM X-gluc and incubated at 37°C in the dark for 3h.

The roots were imaged with a 10x objective lens attached with AmScope T490 Series Simul-Focal Biological Trinocular Compound Microscope and images were captured using a 18MP USB 3.0 C-mount Camera and AmScope software.

### Microscopy

Root hair phenotype in Arabidopsis was observed using a 4x objective lens attached with AmScope T490 Series Simul-Focal Biological Trinocular Compound Microscope and images were captured using a 18MP USB 3.0 C-mount Camera and AmScope software. Maize and wheat root hair phenotypes were images with AmScope SM-1 Series Digital Zoom Trinocular Stereo Microscope 3.5X-225X Magnification on Pillar Stand with Gooseneck LED Lights + 18MP USB 3.0 C-mount USB Camera and AmScope software.

EXP7::NTF and LTI6b-EGFP images were acquired using an Olympus (Evident; Shinjuku, Japan) IXplore SpinSR system, equipped with an inverted IX83 microscope (Evident), a CSUW1-Sora spinning disk confocal unit (Yokogawa) and a Hamamatsu (Shizuoka, Japan) ORCA Fusion BT camera. Fluorescence images were obtained with 10x (maize) or 20x (wheat) objectives. Fluorescence confocal images were acquired using software cellSens (Evident).

### Image analysis

Root hair density and epidermal cell length analysis were performed using ImageJ/Fiji software (Schindelin et al. 2012).

### Statistics

The raw data for each quantification was imported to statistical software JMP Pro17 for generating graphs and performing statistical tests.

## Acknowledgements

Authors thank Miki Fujita and EunKyoung Lee of UBC Bioimaging Facility (RRID: SCR_021304) for their kind support.

## Authors contributions

J.Z., T.D., and M.A.A. designed and performed experiments and analyzed data. J.Z., T.D., and M.A.A. wrote the manuscript and all the authors agreed with the final version of the manuscript.

